# Sacubitril/valsartan (LCZ696) Significantly Reduces Aldosterone and Increases cGMP Circulating Levels in a Canine Model of RAAS Activation

**DOI:** 10.1101/435560

**Authors:** Jonathan P Mochel, Chi Hse Teng, Mathieu Peyrou, Jerome Giraudel, Meindert Danhof, Dean F Rigel

**Author notes:** **Correspondence:** Jonathan P. Mochel, DVM, MS, Ph.D, DECVPT, Associate Professor of Pharmacology, European Veterinary Specialist in Pharmacology and Toxicology, Iowa State University College of Vet. Medicine, 2448 Lloyd, 1809 S Riverside Dr. Ames, IA 50011-1250, Email correspondence.

## Abstract

Simultaneous blockade of angiotensin receptors and enhancement of natriuretic peptides (NP) by the first-in-class angiotensin receptor neprilysin (NEP) inhibitor sacubitril/valsartan constitutes an effective approach to treating heart failure. This study examined the effects of sacubitril/valsartan (225 and 675mg/day) vs. placebo, sacubitril (360mg/day), valsartan (900mg/day), and benazepril (5mg/day) on the dynamics of the renin-angiotensin-aldosterone system (RAAS) and the NP system in dogs. Beagle dogs (n=18) were fed a low-salt diet (0.05% Na) for 15 days to model RAAS activation observed in clinical heart failure. Drugs were administered once daily during the last 10 days, while the effects on the RAAS and NPs were assessed on days 1, 5, and 10. Steady-state pharmacokinetics of the test agents were evaluated on day 5. Compared with placebo, sacubitril/valsartan (675mg) substantially increased cGMP circulating levels, while benazepril and valsartan showed no effect. Additionally, sacubitril/valsartan (675mg) and valsartan significantly increased plasma renin activity, angiotensin I and angiotensin II concentrations. Finally, sacubitril/valsartan (both doses), and valsartan significantly decreased plasma aldosterone vs. placebo. Systemic exposure to valsartan following sacubitril/valsartan 675mg administration was similar to that observed with valsartan 900mg administration alone. Sacubitril/valsartan favorably modulates the dynamics of the renin and NP cascades through complementary NEP and RAAS inhibition.

## 1 Introduction

Chronic heart failure (HF) affects approximately 1–2% of the adult human population in developed countries, with the prevalence rising to ≥10% among persons 70 years of age or older [1]. Importantly, the prognosis of chronic HF remains poor, even with effective adherence to evidence-based pharmacological and non-pharmacological interventions [2, 3] emphasizing the need for novel treatment strategies.

The renin-angiotensin-aldosterone system (RAAS) and natriuretic peptide (NP) cascade are key counterregulatory mechanisms that play a critical role in cardiovascular (CV) physiology and disease pathophysiology. Dysregulation of the RAAS leads to hemodynamic perturbations and end-organ remodeling. Angiotensins I (Ang I) and II (Ang II) mediate vasoconstriction, increase in blood pressure and sympathetic tone, sodium and water retention, aldosterone release, fibrosis, and hypertrophy [4–6]. Furthermore, recent evidence shows that elevated aldosterone levels are associated with reduced survival in patients with hypertension and CV diseases [7], and are a significant prognostic marker in patients with systolic HF [8–10]. In contrast, the NP system inhibits the RAAS and decreases sympathetic activation through cyclic guanosine monophosphate (cGMP)-dependent pathways [11, 12]. Activation of NP receptors increases diuresis and natriuresis, decreases systemic vascular resistance, and plays a protective role in the CV system by counteracting the effects of fluid overload, as well as through anti-proliferative, anti-hypertrophic, and anti-fibrotic mechanisms [11]. Advanced HF constitutes a state of NP deficiency [13] and is associated with a prolonged activation of the RAAS [14, 15]. Although neprilysin (NEP) inhibitors are capable of enhancing NP levels, they are in practice ineffective at lowering blood pressure in hypertensive patients [16], probably due to a concomitant increase in vasoconstrictors such as Ang II and endothelin (ET)-1 [17]. Therefore, simultaneous NEP and RAAS inhibition offers a promising and innovative therapeutic approach in the management of HF. Previous literature showed that concomitant angiotensin converting enzyme (ACE) and NEP inhibition with omapatrilat tended to improve morbidity and mortality in chronic HF but failed to demonstrate substantial benefit over enalapril alone [18]. In addition, omapatrilat was withdrawn from development due to an unacceptably high rate of angioedema in clinical trials [18, 19]. As clinical studies of sacubitril/valsartan (LCZ696) have shown, replacing the ACE inhibitor with an angiotensin receptor blocker (ARB) minimizes the risk of life-threatening angioedema while retaining the beneficial effects of combined NEP and RAAS inhibition [20].

Sacubitril/valsartan is a first-in-class angiotensin receptor neprilysin inhibitor (ARNI), which upon oral administration delivers systemic exposure to sacubitril (AHU377) and valsartan, a well-established ARB recommended by established guidelines for the treatment of HF [1, 21, 22]. Sacubitril is an inactive prodrug that is rapidly hydrolyzed by carboxyl esterase 1 to sacubitrilat, a pharmacologically active NEP inhibitor [23]. Phase II/III clinical trials with sacubitril/valsartan have shown beneficial effects in patients with HF and reduced (HFrEF) or preserved (HFpEF) ejection fraction [20, 24]. Sacubitril/valsartan has been approved in many countries for the treatment of HFrEF and is recommended by European and American HF guidelines [25, 26] for the treatment of chronic symptomatic HFrEF (New York Heart Association Class II–IV).

Similar to humans, activation of the RAAS accompanies the reduced cardiac output reported in canine chronic HF [27, 28], which motivated the choice of this animal species in the present study [29, 30]. Likewise, ACE inhibitors are the standard of care for the treatment of canine chronic HF, and the effects of the ACE inhibitor prodrug benazepril on the renin cascade have been investigated following low-salt-diet activation of the RAAS in dogs [31].

Although numerous studies have shown the positive hemodynamic and clinical effect of sacubitril/valsartan, a comprehensive evaluation of the temporal effects of sacubitril/valsartan on the dynamics of the RAAS and NP cascade is currently missing. The objective of this pharmacology study was to report the pharmacodynamic effects of sacubitril/valsartan on the renin-angiotensin system and cGMP in beagle dogs using a non-invasive model of RAAS activation.

## 2 Methods

### 2.1 Animals

Beagle dogs (N = 18; 9 males and 9 females) from the Novartis Centre de Recherche Sante Animale Test Facility colony (St-Aubin, Switzerland), aged 4–5 years, weighing 10–20 kg, and that were deemed healthy by the study veterinarian, were included in the study.

Suitability for inclusion was evaluated by a physical examination and confirmed by measuring selected hematological (red and white blood cells counts, hemoglobin, hematocrit) and clinical chemistry (albumin, total protein, alanine aminotransferase, aspartate aminotransferase, blood urea nitrogen, creatinine) parameters in blood. Prior to the study start, dogs were acclimatized to the experimental facility for a week. Animals were housed in pens (about 2 m2/animal) containing granulate bedding material and an additional elevated platform for resting. The study rooms had natural daylight and additional artificial light of similar intensity (400 lux) from 07:00 to 19:00 h. Room temperature and relative humidity were within the target ranges of 17–23°C and 35–75%, respectively. The quality of drinking water was compliant with the Swiss Federal Regulations on Foodstuff and was offered *ad libitum*.

### 2.2 Sample size

In absence of preliminary data on the effect of sacubitril/valsartan on biomarkers of the RAAS and cGMP in dogs, a formal sample size calculation with predefined power and type 1 error could not be performed. Instead, determination of the study sample size was based on data evaluation of mean differences in plasma renin activity (PRA) between standard of care benazepril and placebo from a previous experiment in 12 beagle dogs (N = 6 per group, [32], for a type 1 error α = 0.05 with statistical power = 0.80.

### 2.3 Experimental model

The non-invasive and fully reversible low-sodium diet (0.05% Na) was used to model activation of the RAAS, as observed in the course of HF. The low-sodium diet has been established as a reliable and reproducible model of RAAS activation to evaluate the effect of RAAS inhibitors in multiple studies [6, 32, 33]. The experimental procedures were performed in compliance with the registered permit number 10/09 covering animal experiments for CV research in dogs, adopted and approved by the Cantonal Animal Welfare Committee of Fribourg (Switzerland): ‘*Modèle de régime hyposodé pour les maladies cardiovasculaires chez le chien*’ in February 2009. The study protocol was designed to use the fewest number of animals possible while being consistent with the scientific needs of the study, and conformed to international ethical standards [34].

### 2.4 Study design

A 3-way partial crossover study design was chosen to examine the effect of the test and reference treatment items over a period of 10 days (Figure 1). To achieve steady-state activation of RAAS biomarkers, animals were fed a low-sodium diet for 5 days prior to the oral administration of the study drugs: sacubitril (SAC) calcium salt at 360 mg (Period A); a low sacubitril/valsartan tri-sodium hemipentahydrate salt dose (SVL) at 225 mg (period A); a high sacubitril/valsartan dose (same formulation) (SVH) at 675 mg (Period B and C); valsartan free acid (VAL) 900 mg (Period B and C); benazepril hydrochloride (BNZ) 5 mg (Period C); and empty capsules as placebo (PBO; period A and B) (Figure 1). Dogs were administered the appropriate treatment at 7:00 AM and were fed 12 hours thereafter following withdrawal of the +12-hour blood sample. A 2-week washout period with the dogs on normal chow was maintained between each successive treatment Period.

**Figure 1.**
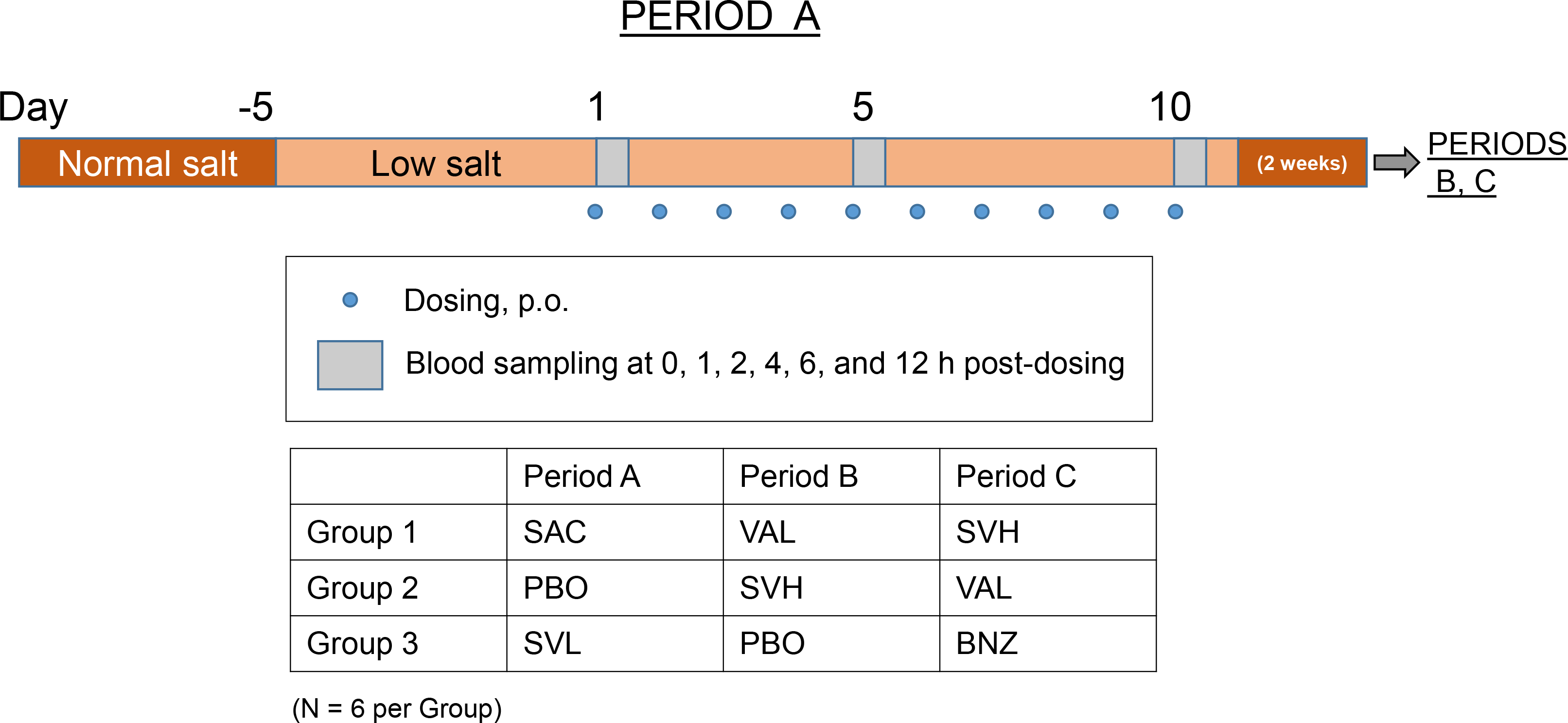
A 3-way partial crossover study design was chosen to examine the effect of sacubitril/valsartan over 10 days. To achieve steady-state activation of RAAS biomarkers, animals were fed a low-salt diet for 5 days prior to the oral administration of the study drugs: sacubitril (SAC) calcium salt at 360 mg (Period A); a low sacubitril/valsartan tri-sodium hemipentahydrate salt dose (SVL) at 225 mg (period A); a high sacubitril/valsartan dose (same formulation) (SVH) at 675 mg (Period B and C); valsartan free acid (VAL) 900 mg (Periods B and C); benazepril hydrochloride (BNZ) 5 mg (Period C); and empty capsules as placebo (PBO; Periods A and B). Low-salt diet was continued throughout the 10 treatment days. Two weeks of washout with the dogs on normal chow were incorporated between each successive treatment Period.

### 2.5. Drug dose justification

The selected nominal doses of the therapeutic drugs corresponded to SAC ~24 mg/kg, SVL and SVH ~15 and ~45 mg/kg, VAL ~60 mg/kg, and BNZ ~0.33 mg/kg for an average 15 kg body weight dog. The SVL and SVH doses were selected based on internal preliminary efficacy/safety evaluation and drug metabolism and pharmacokinetics/safety studies in dogs. The dose of VAL was selected to match the exposure of valsartan from the SVH group. This was based on previous findings from Gu et al. [35], showing that the oral bioavailability of VAL following LCZ696 administration was about 3-fold higher than that observed following administration of approximately equimolar doses of VAL alone. The dose of SAC was chosen as an approximately equimolar dose to the NEP inhibitor delivered by the SVH dose. The 5 mg BNZ dose corresponds to the recommended dose in dogs with chronic HF [36].

### 2.5. Pharmacodynamic assessment

Due to the known sensitivity of the renin-angiotensin cascade to posture and external stimuli [37], specific precautions were taken: i) dogs were maintained in a standing position during blood collection, ii) sampling was performed in a sound-protected room, and iii) low-intensity lighting was used during withdrawal. Blood samples were collected from the *vena jugularis* or exceptionally from the vena *cephalixa antebrachii* into 1.2 or 2.7 mL S-Monovette^®^ tubes (Sarstedt Inc. Newton, NC, USA) and kept on ice until centrifugation under refrigeration (2 ± 1°C), as described by [6, 38, 39]. Samples were collected at pre-dose, and at 1, 2, 4, 6, and 12 hours after oral administration on dosing days 1, 5, and 10 for pharmacodynamic assessment of PRA, plasma angiotensin I (Ang I) and II (Ang II), ALD, and plasma cGMP (Periods B and C only). Plasma samples for cGMP determination were not collected during period A and results are therefore not available for SAC and SVL. Measurements of plasma ALD concentrations were carried out using validated high-performance liquid chromatography-mass spectrometry (LC-MS/MS) method with a lower limit of quantification (LLOQ) of 0.02 ng/mL. Plasma cGMP (enzyme immunoassay [EIA] kit, Cayman Chemical Company, USA), Ang I (liquid solid extraction kit, Bachem S-1188, Switzerland), Ang II (EIA kit, SPI BIO, France), and PRA (EIA kit, USCN Life Sciences Inc, China) were performed using validated kits, as previously described. PRA was determined by measuring the rate of Ang I formation after 2-hour incubation of endogenous renin and angiotensinogen in plasma at 37°C and pH 7.2. The LLOQs were 30 pg/mL and 2 pg/mL for Ang I and Ang II, respectively, and 0.05 pmol/mL for cGMP. Analyses were performed in duplicates; values with a coefficient of variation below 25% were retained for statistical evaluation.

### 2.6. Pharmacokinetic assessment

Pharmacokinetic measures were performed using blood collected at day 5 from the *vena jugularis* (or exceptionally from the *vena cephalica antebrachii*) into 1.2 mL S-Monovette^®^ tubes (Sarstedt Inc. Newton, NC, USA). For sample collection of BNZ, heparin was used as an anticoagulant, and for VAL, SVL/SVH, and SAC, EDTA was used. The tubes were gently inverted 5 times and chilled in ice immediately, then centrifuged at 1600 g for 15 minutes at 1°C to obtain the plasma specimen. Plasma samples were frozen at –80°C until further analysis. Plasma concentrations of sacubitrilat, benazeprilat, and VAL were analyzed in the SAC-, SVL/SVH-, BNZ-, and VAL-treated dogs. Concentrations of sacubitrilat, benazeprilat, and VAL were determined using validated high-performance LC-MS/MS methods. The LLOQ of benazeprilat in plasma was 0.5 ng/mL, and 5 ng/mL for sacubitrilat and VAL.

Pharmacokinetic parameter estimates were derived from a statistical moment (non-compartmental) analysis implemented in validated SAS macros (SAS^®^ Version 9.1) and consisted of the following:

1. the maximum concentration (C_max_),
2. the time to maximum concentration (T_max_), and
3. the area under the concentration‒time curve (AUC_0-last_).

Pre-dose time was specified with time 0 (hour) and corresponding values below LLOQ were replaced by zero. Below LLOQ values at subsequent times were excluded from the analysis. Summary statistics including geometric mean and range of values were provided for all mentioned pharmacokinetic parameters.

### 2.7. Safety evaluation

Safety assessments included hematology, biochemistry, hemostasis, body weight, and body condition scoring.

### 2.8. Statistical analyses of biomarker data

To anticipate plausible variations in biomarker levels across treatment days, data were expressed as absolute change from baseline, defined as the individual biomarker concentration at hour 0 (pre-dose), separately for day 1, day 5 and day 10. In accordance with previous descriptions of the effect of sacubitril/valsartan on the RAAS in humans [35], individual time-weighted average (TWA) change from baselines were estimated separately for each day (D1, D5 and D10) in each period (A, B, C), and analyzed by random effect-repeated measures analyses of variance (RRMANOVA), with fixed effect classification variables PERIOD (A, B and C), TRT (treatment group with 6 levels: Placebo, SAC, SVL, SVH, VAL, and BNZ), DAY (D1, D5, and D10) and the two-way interaction TRT by DAY. ANIMAL (1 to 18) was included as a random effect in the model.

Finally, in order to leverage all available pharmacodynamic information and derive meaningful and robust statistical comparisons, plasma biomarker data (both time courses and TWAs) from days 1, 5, and 10 were pooled for each treatment and analyzed by the RRMANOVA approach. All calculations were done using SAS^®^ Version 9.2 by applying univariate analysis for calculation of summary statistics. SAS^®^ procedure was applied to execute the analyses of variance. All tests were performed two-sided with a level of significance α pre-defined at 0.05.

## 3 Results

All experimental animals were randomly assigned to three groups of 6 dogs each and were available for pharmacodynamic, pharmacokinetic, and safety assessments. No statistical difference in baseline characteristics were observed between study groups, based on selected hematological and clinical chemistry parameters.

### 3.1. Safety assessment

All experimental animals completed the study without any incidence of adverse events with any of the test drugs. All dogs returned to the maintenance facility at the end of the experiment.

### 3.2. Effect on plasma cGMP

The typical baseline value for cGMP across treatment groups was ca. 15 pmol/mL. RRMANOVA results showed significant increases in cGMP circulating levels within all treatment groups, including PBO (Figure 2, Panels A and B), which are indicative of diurnal variations of this biomarker in dogs. The TRT effect was found to be significant, but the TRT by DAY interaction did not reach the level of statistical significance. The estimated differences in TWA changes in cGMP were significant between the SVH group and the other three treatment groups (Figure 2, Panel C). On an average SVH significantly increased circulating cGMP levels by approximately 4 pmol/mL as compared with VAL, BNZ, and PBO (Figure 2, Panel C). Conversely, no apparent differences were reported between VAL, BNZ, and PBO treated dogs.

**Figure 2.**
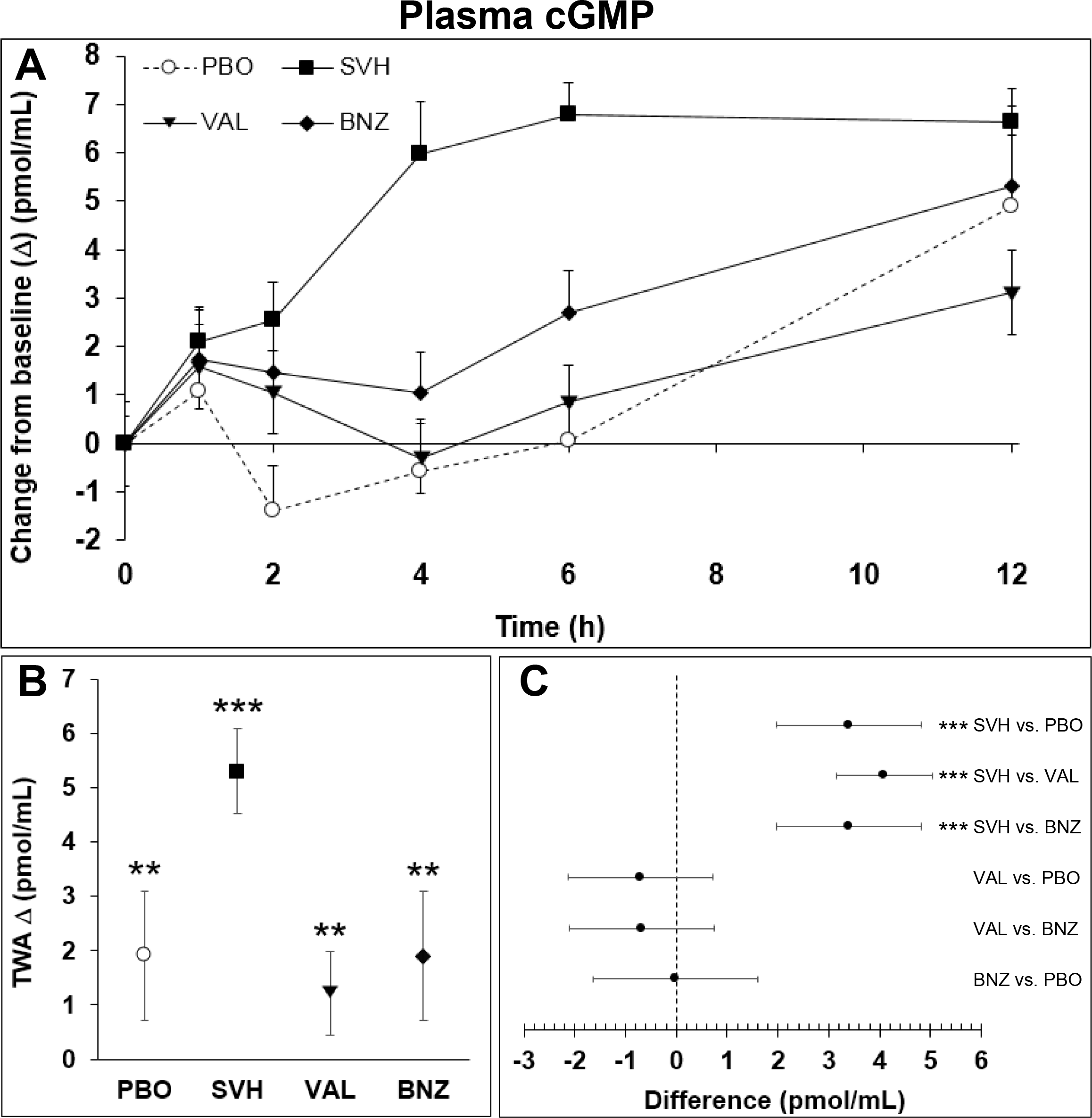
Pharmacodynamics of sacubitril/valsartan (SVH: 675 mg) compared with standard of care benazepril (BNZ: 5mg), sacubitril (SAC: 360 mg) and valsartan (VAL: 900 mg) alone on plasma cGMP. (A) Temporal (absolute) change from baseline (pmol/mL): mean ± S.E.M; (B) Time-weighted average (TWA, pmol/mL) change from baseline (∆): mean + 95% CI; (C) Between-group differences: mean + 95% CI. *0.01 ≤ p < 0.05; **: 0.001 ≤ p < 0.01; ***: p < 0.001.

### 3.3. Effect on PRA

The typical baseline value for PRA across treatment groups was ca. 400 pg/mL/h. PRA remained relatively stable over the 12-hour observation period in the PBO and SAC groups, but appeared to increase with the remaining treatments, and especially with SVH and VAL (Figure 3, Panel A). Results from the RRMANOVA showed that only the TRT effect was significant. The PERIOD and DAY effect, and the TRT by DAY interaction were not statistically significant. The effect of sacubitril/valsartan on PRA was dose-dependent, with only SVH and VAL showing a significant TWA change from baseline (Figure 3, Panel B). Likewise, both SVH and VAL achieved significantly greater elevation of PRA than PBO, SAC, and BNZ (Figure 3, Panel C). Also, VAL showed significantly greater increase in PRA than SVL and SVH (Figure 3, Panel C).

**Figure 3.**
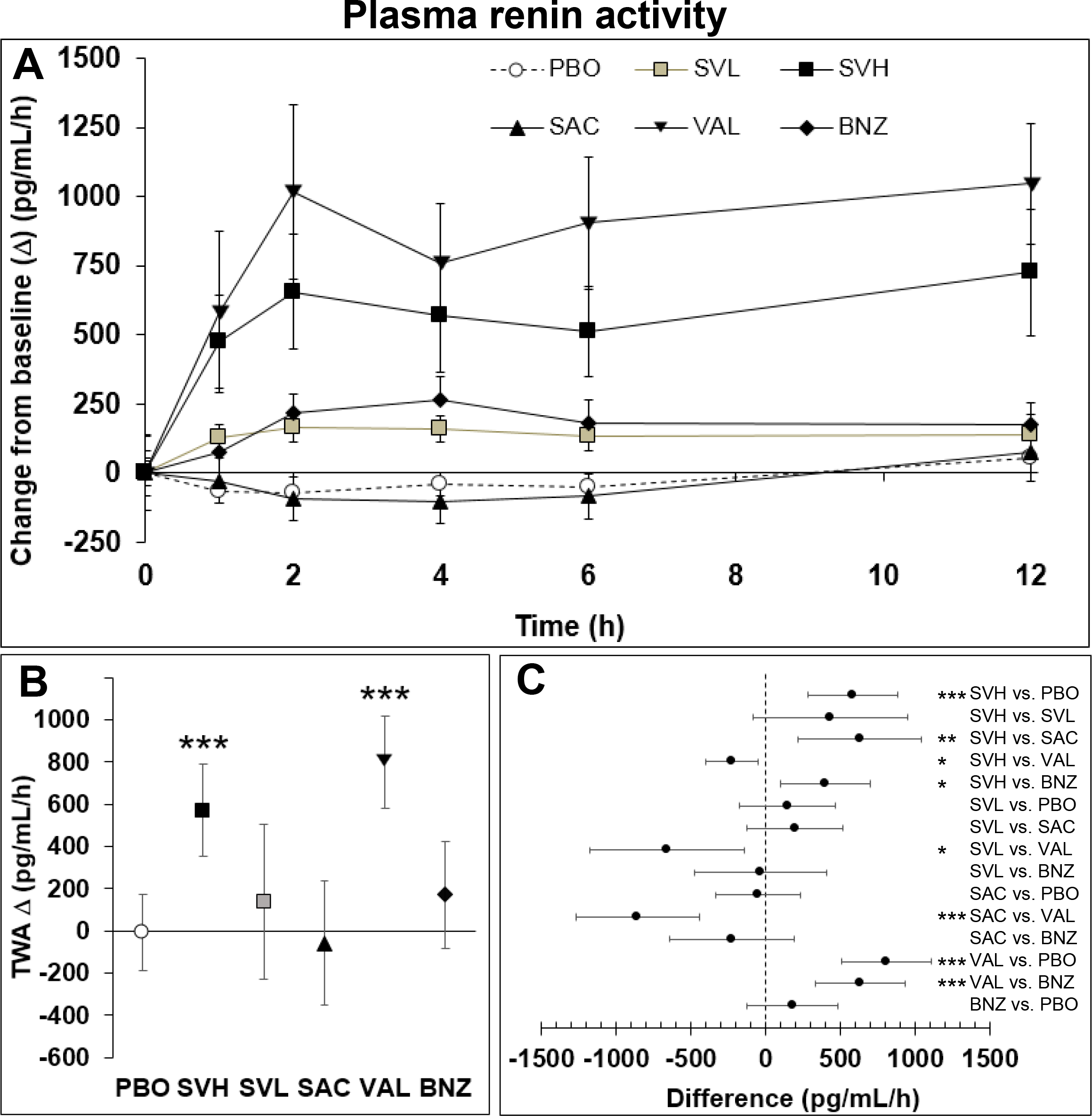
Pharmacodynamics of sacubitril/valsartan (SVL: 225 mg; SVH: 675 mg) compared with standard of care benazepril (BNZ: 5mg), sacubitril (SAC: 360 mg) and valsartan (VAL: 900 mg) alone on plasma renin activity. (A) Temporal (absolute) change from baseline (pg/mL/h): mean ± S.E.M; (B) Time-weighted average (TWA, pg/mL/h) change from baseline (∆): mean + 95% CI; (C) Between-group differences: mean + 95% CI. *0.01 ≤ p < 0.05; **: 0.001 ≤ p < 0.01; ***: p < 0.001.

### 3.4. Effect on Ang I and Ang II

The time-course of response for Ang I and Ang II appeared seemingly consistent with that of PRA. The typical baseline value under low-sodium diet was ca. 185 pg/mL and 15 pg/mL for Ang I and Ang II, respectively. For both angiotensins, the results of the RRMANOVA showed that only the TRT effect was significant. The PERIOD and DAY effect, and the TRT by DAY interaction were not significant.

An apparent increase in Ang I was observed for all treatment groups but SAC and PBO, with VAL and SVH showing the most pronounced effect overall (Figure 4, Panel A). Similar to PRA, only SVH and VAL showed a significant TWA Ang I change from baseline (Figure 4, Panel B). Differences to PBO were highly significant for both sacubitril/valsartan dosing groups. The effect of VAL was superior to that of all other treatment groups, while dosing with SVH and BNZ yielded significant differences to sacubitril alone (Figure 4, Panel C).

**Figure 4.**
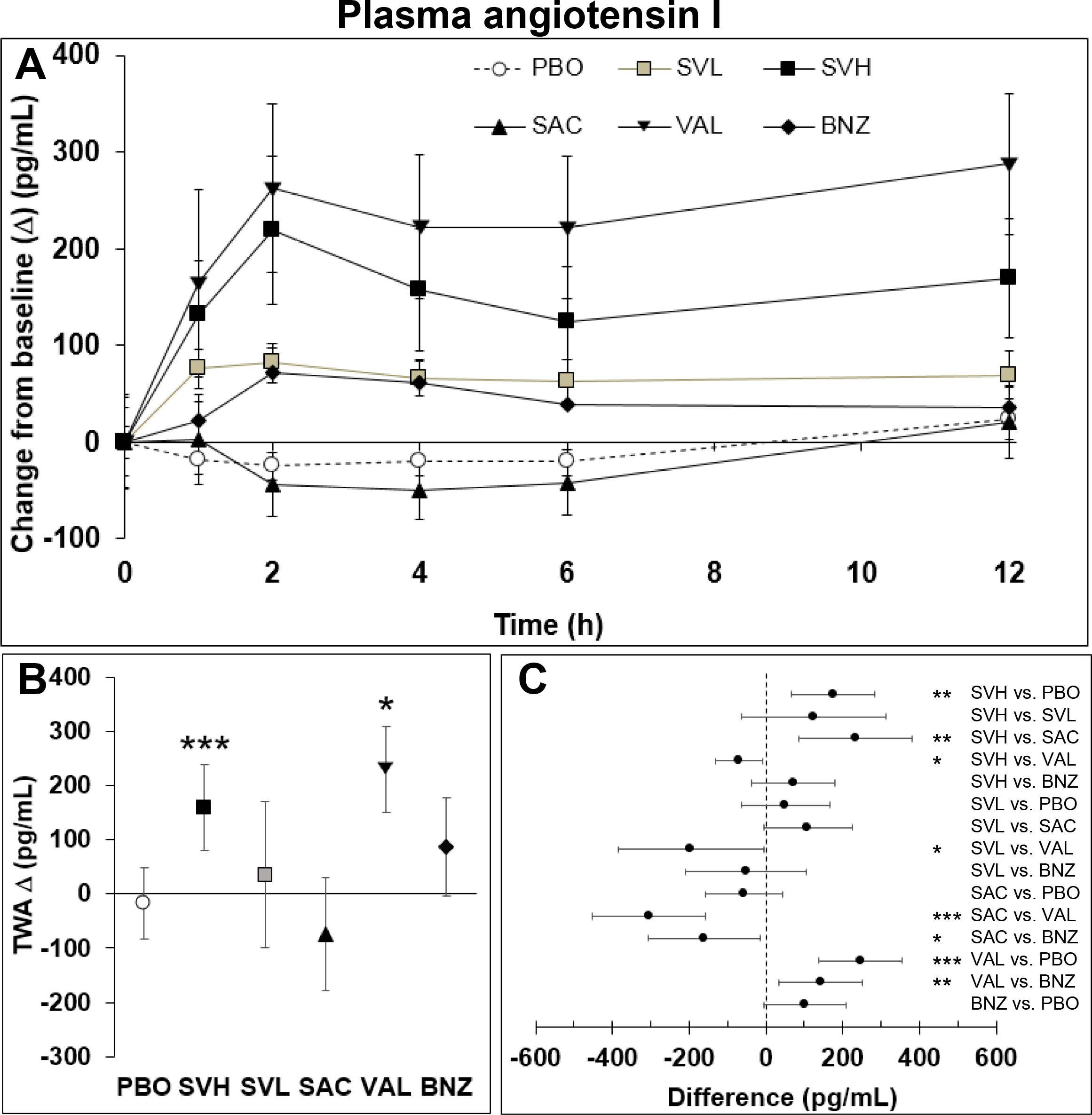
Pharmacodynamics of sacubitril/valsartan (SVL: 225 mg; SVH: 675 mg) compared with standard of care benazepril (BNZ: 5mg), sacubitril (SAC: 360 mg) and valsartan (VAL: 900 mg) alone on plasma angiotensin I. (A) Temporal (absolute) change from baseline (pg/mL): mean ± S.E.M; (B) Time-weighted average (TWA, pg/mL) change from baseline (∆): mean + 95% CI; (C) Between-group differences: mean + 95% CI. *0.01 ≤ p < 0.05; **: 0.001 ≤ p < 0.01; ***: p < 0.001.

All treatment groups but PBO and BNZ appeared to elevate Ang II, with sacubitril/valsartan showing the most pronounced effect overall (Figure 5, Panel A). Consistent with PRA and Ang I, only SVH and VAL demonstrated a significant TWA Ang II change from baseline (Figure 5, Panel B). SVH and VAL significantly increased Ang II as compared with PBO (estimated difference of 9.6 and 6.7 pg/mL, respectively) (Figure 5, Panel C). There was a modest and non-significant increase in Ang II following SVL (+4.4 pg/mL vs. PBO), and SAC treatment alone (+2.8 pg/mL vs. PBO).

**Figure 5.**
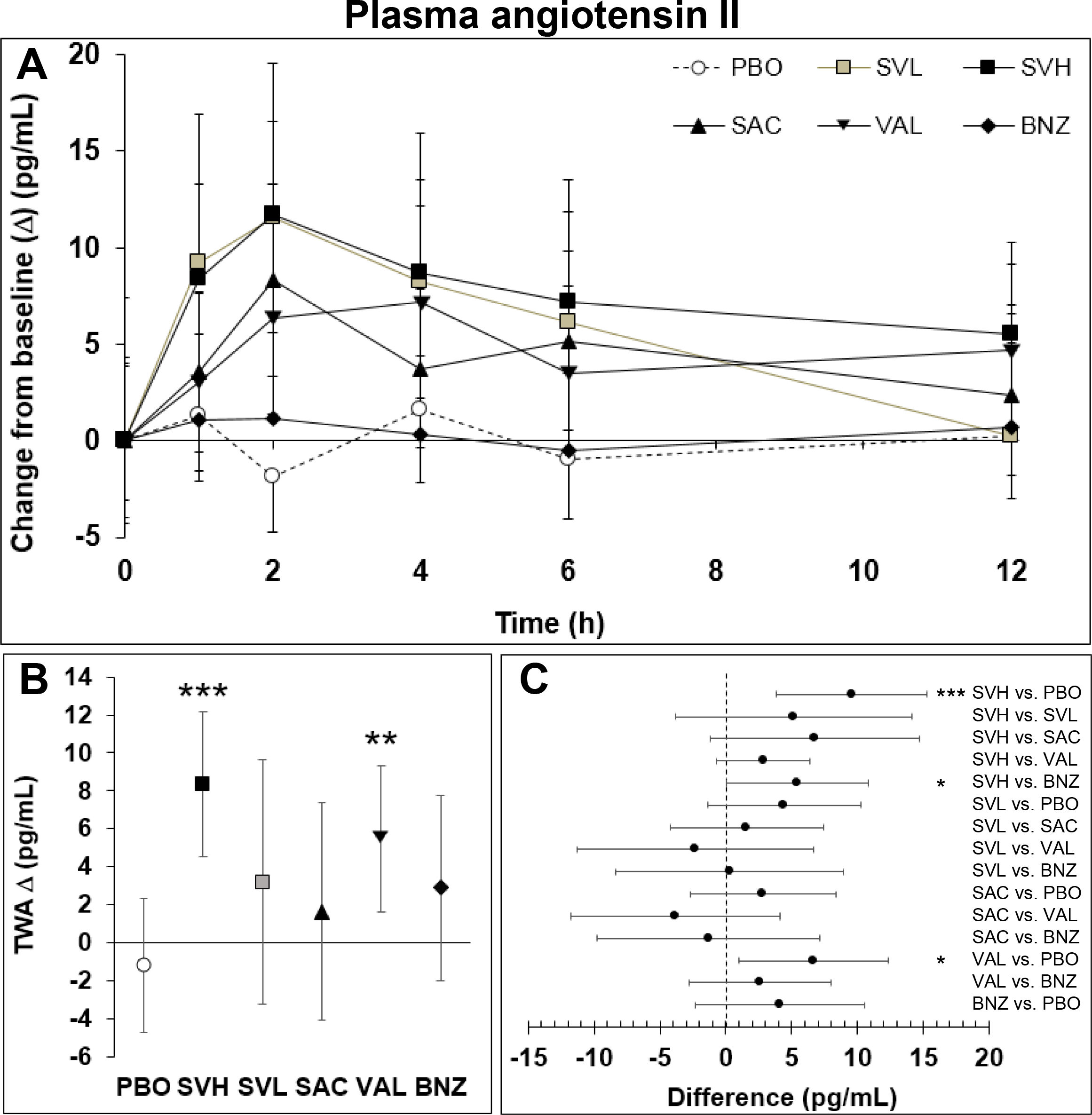
Pharmacodynamics of sacubitril/valsartan (SVL: 225 mg; SVH: 675 mg) compared with standard of care benazepril (BNZ: 5mg), sacubitril (SAC: 360 mg) and valsartan (VAL: 900 mg) alone on plasma angiotensin II. (A) Temporal (absolute) change from baseline (pg/mL): mean ± S.E.M; (B) Time-weighted average (TWA, pg/mL) change from baseline (∆): mean + 95% CI; (C) Between-group differences: mean + 95% CI. *0.01 ≤ p < 0.05; **: 0.001 ≤ p < 0.01; ***: p < 0.001.

### 3.5. Effect on ALD

The typical baseline value for ALD under low-sodium diet was ca. 0.25 ng/mL. Both sacubitril/valsartan doses and VAL had an apparent effect on ALD plasma concentrations, while only modest and non-significant changes were reported in the other treatment groups (Figure 6, Panel A). Interestingly enough, the decrease of ALD in the sacubitril/valsartan groups was not dose-dependent, and a rebound of ALD concentration was observed in the SVH dosing group. Results from the RRMANOVA showed that only the TRT effect was significant. The PERIOD and DAY effect, and the TRT by DAY interaction were not of significance. SVH, SVL VAL and BNZ achieved a significant TWA change from baseline (Figure 6, Panel B). In addition, differences to PBO were found to be statistically significant for SVH, SVL and VAL, but not significant for BNZ (Figure 6, Panel C). There was a trend towards a decrease of ALD with SAC, but the estimated difference to PBO (approximately half of the reduction obtained with VAL) did not reach the level of statistical significance.

**Figure 6.**
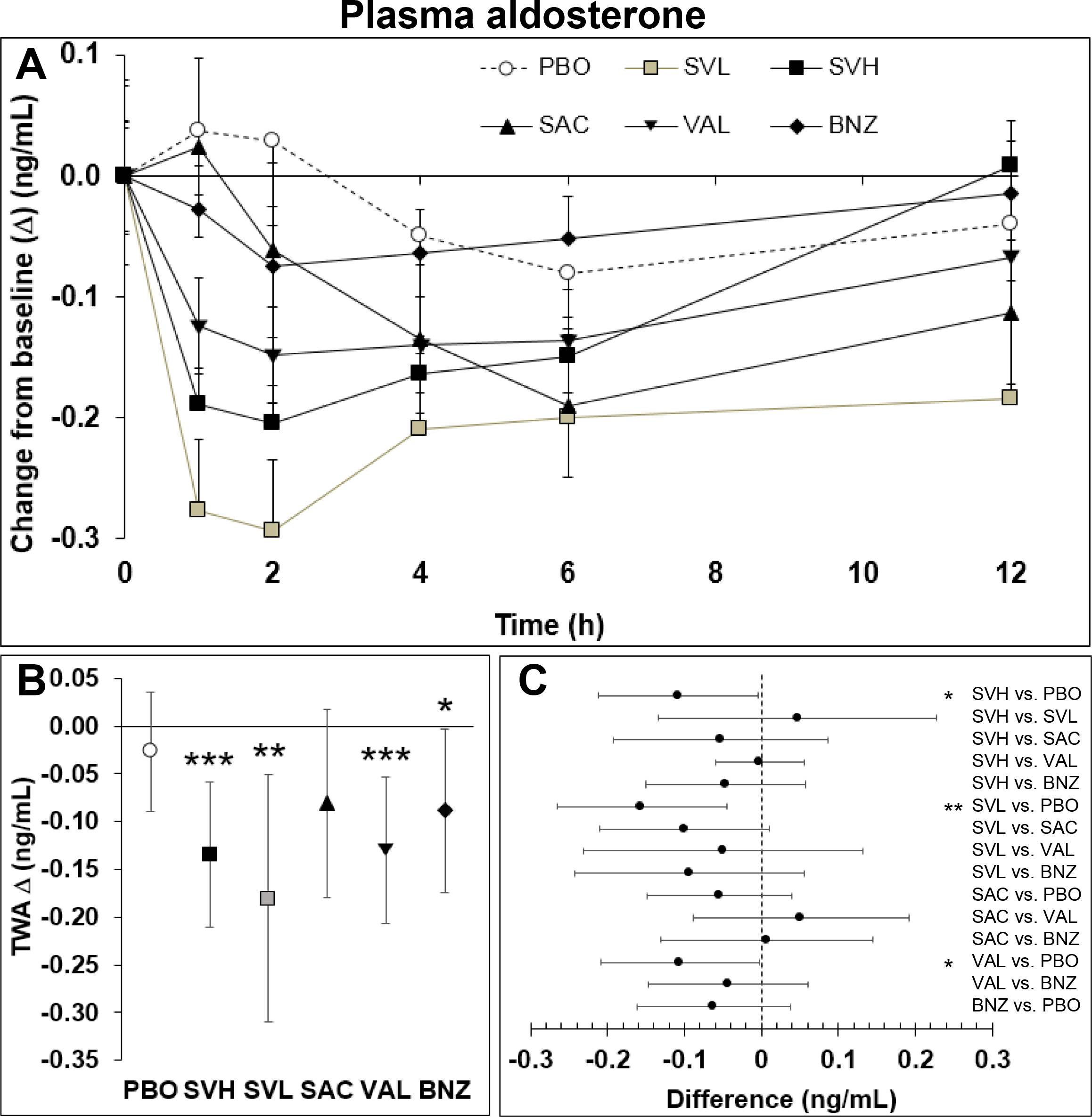
Pharmacodynamics of sacubitril/valsartan (SVL: 225 mg; SVH: 675 mg) compared with standard of care benazepril (BNZ: 5mg), sacubitril (SAC: 360 mg) and valsartan (VAL: 900 mg) alone on plasma aldosterone. (A) Temporal (absolute) change from baseline (ng/mL): mean ± S.E.M; (B) Time-weighted average (TWA, ng/mL) change from baseline (∆): mean + 95% CI; (C) Between-group differences: mean + 95% CI. *0.01 ≤ p < 0.05; **: 0.001 ≤ p < 0.01; ***: p < 0.001.

### 3.6. Pharmacokinetics of the test drugs

Plasma pharmacokinetics following oral dosing with sacubitril/valsartan, SAC, VAL, and BNZ on Day 5 is presented in Table 1. SVH delivered comparable systemic exposure (as defined by C_max_, AUC_0-last_) of sacubitrilat as the SAC 360 mg dose. Systemic exposure to VAL was also similar between SVH and VAL 900 mg. Furthermore, there was an apparent more than dose-proportional increase in exposure to sacubitrilat between the SVL (225 mg) and SVH (675 mg) doses. However, the exposure increase of VAL was approximately proportional with the dose between SVL and SVH. The T_max_ values of the sacubitril/valsartan analytes were similar for both SVL and SVH. The time to maximum sacubitrilat and VAL peak concentrations appeared to be slightly shorter in the sacubitril/valsartan groups as compared with the SAC and VAL alone treatments.

**Table 1.** Pharmacokinetic parameters on Day 5 in beagle dogs on a low salt diet.

|  |  | **Sacubitrilat** | **Valsartan** | **Benazeprilat** |
| --- | --- | --- | --- | --- |
|  | **Group (Dose)** | **Mean (± S.D)** | **Mean (± S.D)** | **Mean (± S.D)** |
| **C<sub>max</sub>** (ng/mL) | SAC (360 mg) | 2686 (± 2734) | — | — |
|  | SVL (225 mg) | 348 (± 150) | 917 (± 486) | — |
|  | SVH (675 mg) | 2103 (± 2461) | 2288 (± 1582) | — |
|  | VAL (900 mg) | — | 2769 (± 2178) | — |
|  | BNZ (5 mg) | — | — | 24 (± 8) |
| **T<sub>max</sub>** (hour) | SAC (360 mg) | 3.0 (± 1.0) | — | — |
|  | SVL (225 mg) | 1.7 (± 1.0) | 2.5 (± 2.0) | — |
|  | SVH (675 mg) | 2.0 (± 0.6) | 2.8 (± 3.1) | — |
|  | VAL (900 mg) | — | 4.2 (± 3.8) | — |
|  | BNZ (5 mg) | — | — | 2.2 (± 1.2) |
| **AUC<sub>0-last</sub>** (ng/mL*h) | SAC (360 mg) | 9026 (9289) | — | — |
|  | SVL (225 mg) | 1062 (295) | 4325 (± 1537) | — |
|  | SVH (675 mg) | 6244 (6311) | 14483 (± 9363) | — |
|  | VAL (900 mg) | — | 17396 (± 11347) | — |
|  | BNZ (5 mg) | — | — | 132 (± 40) |

## 4 Discussion

We report the results of the first comprehensive evaluation of the temporal effects of sacubitril/valsartan on biomarkers of the RAAS and cGMP using an established canine model of RAAS activation.

Consistent with previous findings from Gu et al., [35], our pharmacokinetic analysis showed that systemic exposure to VAL (AUC_0-last_ and C_max_) following sacubitril/valsartan oral administration was about 3-fold higher than that observed after approximately equimolar doses of VAL. Consequently, the exposure to VAL between the 900 mg VAL group and the 675 mg sacubitril/valsartan oral treatments was comparable, such that differences in RAAS and cGMP biomarkers between these two groups can be attributed to NEP inhibition alone. The large between-dog variation in VAL and sacubitrilat exposure was expected and is in agreement with previous results from Gu et al. [35] who reported a coefficient of variation between 50% and 100% in a preliminary pharmacokinetic study with 3 Beagle dogs. At this time, the structural causes of such variability are unclear, yet it had apparent consequences on the variations of the RAAS and cGMP biomarkers response to SVH, SVL and VAL, limiting statistical comparisons between study groups. Finally, the pharmacokinetics of benazeprilat following 5 mg oral dosing with BNZ is consistent with previous literature in dogs [36, 40].

Inhibition of NEP is known to be associated with increased levels of NPs, which stimulate synthesis of cGMP [35]. The observed trend towards plasma cGMP increase in the late afternoon for all treatment groups is consistent with published literature in humans [41], which is indicative of diurnal oscillations of this biomarker in dogs. While sacubitril/valsartan, VAL, and BNZ modulated the renin cascade, the increased cGMP levels by sacubitril/valsartan, but not VAL or BNZ, demonstrate activation of the NP system attributable to sacubitrilat, the active metabolite of SAC [35, 42]. These changes could also be mediated (at least in part) by variations in circulating nitric oxide (NO) levels as recent studies in rats showed an increase in NO bioavailability consecutive to sacubitril/valsartan treatment [43]. Although cGMP data for the SVL dose were not available in the present study, previous publications in healthy humans have demonstrated dose-dependent increases in circulating cGMP levels. In these experiments, plasma cGMP increased as early as 4 hours following sacubitril/valsartan administration compared with placebo, with a return to baseline level within 24 hours [35]. Similarly, in the same study, dose dependent increases in RAAS biomarkers (PRA and Ang II) reached maximum within 4 hours of sacubitril/valsartan dose in healthy human participants [35]. In patients with HF and left ventricular ejection fraction ≤40%, sacubitril/valsartan 100 mg titrated to 200 mg twice daily increased plasma cGMP (1.4 times baseline) and urinary ANP as a result of NEP inhibition [44]. Likewise, significant reductions of plasma NT-pro brain NP in patients with HFrEF treated with sacubitril/valsartan 200 mg showed clinical benefits in the PARADIGM-HF study, which correlated with risk reduction in CV mortality and HF hospitalizations compared with the ACE inhibitor enalapril [20, 45].

SVH and VAL alone significantly increased PRA over the course of the study. Similar to PRA, plasma Ang I levels increased in both groups, indicating that sacubitril/valsartan blocks Ang II signaling through the Ang II type 1 (AT_1_) receptor, causing the known compensatory up-regulation of plasma renin and Ang I [46, 47]. In contrast, the ACE inhibitor benazepril did not significantly increase PRA, which was unexpected and in contradiction with previous literature reporting similar ranges of systemic exposure to benazeprilat in dogs [6, 32].

Interestingly, VAL showed significantly greater elevation of PRA than SVL and SVH. Likewise, SAC and SVL did not show a significant effect on the levels of Ang I, while the effect of VAL on Ang I appeared to be significantly greater than the effect of SVH. These observations are consistent with the known effect of atrial NP to suppress renin production [48], thereby leading to a more pronounced effect of VAL on PRA and Ang I as compared with SVH.

Overall, the increase in plasma Ang II levels in response to treatment was similar to the changes observed for PRA and Ang I, except for the effect of SVH being somewhat more pronounced than VAL at the early time-points. This is consistent with the inhibition of NEP, an enzyme known to degrade plasma Ang II [49, 50]. In contrast, BNZ had no noticeable effect on plasma Ang II levels, which is also in line with our findings on PRA and Ang I. This upholds previous reports showing only partial reduction of Ang II in dogs receiving 10 mg BNZ, and no decrease in circulating Ang II in 45% of canine patients with stable chronic HF despite long-term ACE inhibitor use. One possible explanation is activation of alternative biological pathways (e.g. chymase, cathepsin G and tonin) for Ang II production [6, 31, 32, 51].

Both sacubitril/valsartan doses and VAL significantly decreased ALD levels, with the greatest decrease observed in the sacubitril/valsartan treated groups within the first 2 hours after dosing. In addition, SAC showed a moderate (but non-significant) decrease in ALD levels compared with placebo (reaching approximately half of the reduction in ALD observed with VAL), indicating that simultaneous inhibition of NEP and blockade of the AT_1_ receptor by sacubitril/valsartan could in theory be additive and lead to positive clinical outcomes by decreasing a known prognostic marker of HF. Of note, oral dosing with SVH and VAL did result in comparable reduction of ALD in dogs (while providing similar exposure to VAL), which would indicate that the effect of sacubitril/valsartan on ALD is mainly driven by VAL. In humans, the degree of ALD increase is related to the severity of heart failure [52] and ALD is known to worsen Ang II tissue-damaging properties [53]. Therefore, elevated exposure to ALD has been associated with a poor prognosis in multiple case studies [54, 55]. More precisely, Swedberg et al. [56] have found a positive correlation between mortality and systemic levels of ALD (p < 0.003) in a group of severe HF patients. In a report from Güder et al. [8], high ALD concentrations were found to be a predictor of increased mortality risk that provides complementary prognostic value in a prospective cohort experiment of 294 HF patients. Finally, and consistently with our observations in dogs, the clinical relevance of RAAS inhibition and ALD reduction in patients under ARNI therapy was demonstrated in a study by Jordaan et al [57].

Interestingly, a steep ALD return to baseline was noted in the SVH group between 6 and 12 hours after dosing, implicating a rebound phenomenon occurring at the higher sacubitril/valsartan dose and suggesting optimum ALD inhibition being achieved at a lower therapeutic dose. Similar rebound in ALD levels were observed after infusion of atrial NP in patients with mild-to-moderate hypertension [58].

### 4.1. Limitations

Because of the small study size, the statistical significance of certain findings was hampered by low statistical power. This is illustrated by the non-significance of the inhibitory effect of SAC alone on ALD, PRA and Ang I, and its stimulatory effect on Ang II. Conversely, the clinical significance of the reported statistical differences between study groups remains unclear, although, these are consistent with clinical results from the PARADIGM-HF study demonstrating superiority of sacubitril/valsartan over enalapril in human patients with HF. In addition, the effect of low dose sacubitril/valsartan on plasma cGMP could not be investigated in the present study, leaving it unclear whether sufficient inhibition of NEP could be achieved with a 225 mg dose. Finally, results from our earlier research [59] have shown an 8- to 10-fold rise in urinary ALD in 6 healthy beagle dogs fed a low-salt diet (0.05% Na) for 10 days. While sodium restriction is a powerful stimulant of the renin-angiotensin cascade, a detailed description of Ang II and ALD elevation in dogs suffering from HF is currently missing. This would be an important step towards the formal validation of the low-salt diet as a reliable model of HF-related RAAS activation. As such, the positive pharmacological effects of sacubitril/valsartan reported in the present study should be confirmed by additional clinical work in dogs with HF to evaluate the hemodynamic effect of ARNI on disease modulation in canines.

### 4.2. Conclusion

In conclusion, the ARNI sacubitril/valsartan reduced ALD, a known risk factor of CV mortality, and enhanced the NP system via cGMP-mediated pathways in a low-sodium diet model of RAAS activation. The results presented herein provide further evidence that the effects on the renin cascade extend to reduced ALD levels beyond that achieved with RAAS blockade alone. These positive findings in dogs also suggest that sacubitril/valsartan is a promising pharmacological candidate for increased survival in canine cardiovascular diseases.

## 7 Author Contributions

JPM, MP and DFR conceived the experimental protocols. JG and MD contributed to the development of the hypothesis and reviewed the study protocols. CHT was responsible for the statistical analysis of the study results. All authors contributed to the preparation of the manuscript.

## 8 Acknowledgments

All contributing authors had full access to the study data and agree with the publication of the results.

## References

1 McMurray JJ, Adamopoulos S, Anker SD, Auricchio A, Bohm M, Dickstein K, Falk V, Filippatos G, Fonseca C, Gomez-Sanchez MA, Jaarsma T, Kober L, Lip GY, Maggioni AP, Parkhomenko A, Pieske BM, Popescu BA, Ronnevik PK, Rutten FH, Schwitter J, Seferovic P, Stepinska J, Trindade PT, Voors AA, Zannad F, Zeiher A, Task Force for the D, Treatment of A, Chronic Heart Failure of the European Society of C, Bax JJ, Baumgartner H, Ceconi C, Dean V, Deaton C, Fagard R, Funck-Brentano C, Hasdai D, Hoes A, Kirchhof P, Knuuti J, Kolh P, McDonagh T, Moulin C, Popescu BA, Reiner Z, Sechtem U, Sirnes PA, Tendera M, Torbicki A, Vahanian A, Windecker S, McDonagh T, Sechtem U, Bonet LA, Avraamides P, Ben Lamin HA, Brignole M, Coca A, Cowburn P, Dargie H, Elliott P, Flachskampf FA, Guida GF, Hardman S, Iung B, Merkely B, Mueller C, Nanas JN, Nielsen OW, Orn S, Parissis JT, Ponikowski P, Guidelines ESCCfP (2012) ESC guidelines for the diagnosis and treatment of acute and chronic heart failure 2012: The Task Force for the Diagnosis and Treatment of Acute and Chronic Heart Failure 2012 of the European Society of Cardiology. Developed in collaboration with the Heart Failure Association (HFA) of the ESC. Eur J Heart Fail. 14 (8): 803–69.https://doi.org/10.1093/eurjhf/hfs105

2 Azad N, Lemay G (2014) Management of chronic heart failure in the older population. J Geriatr Cardiol. 11 (4): 329–37.https://doi.org/10.11909/j.issn.1671-5411.2014.04.008

3 Oldland E, Driscoll A, Currey J (2014) High complexity chronic heart failure management programmes: programme characteristics and 12 month patient outcomes. Collegian. 21 (4): 319–26

4 Hsueh WA, Wyne K (2011) Renin-Angiotensin-aldosterone system in diabetes and hypertension. J Clin Hypertens (Greenwich). 13 (4): 224–37.https://doi.org/10.1111/j.1751-7176.2011.00449.x

5 Mishra A, Srivastava A, Mittal T, Garg N, Mittal B (2012) Impact of renin-angiotensin-aldosterone system gene polymorphisms on left ventricular dysfunction in coronary artery disease patients. Dis Markers. 32 (1): 33–41.https://doi.org/10.3233/DMA-2012-0858

6 Mochel JP, Peyrou M, Fink M, Strehlau G, Mohamed R, Giraudel JM, Ploeger B, Danhof M (2013) Capturing the dynamics of systemic Renin-Angiotensin-Aldosterone System (RAAS) peptides heightens the understanding of the effect of benazepril in dogs. J Vet Pharmacol Ther. 36 (2): 174–80.https://doi.org/10.1111/j.1365-2885.2012.01406.x

7 Pimenta E, Gordon RD, Stowasser M (2013) Salt, aldosterone and hypertension. J Hum Hypertens. 27 (1): 1–6.https://doi.org/10.1038/jhh.2012.27

8 Guder G, Bauersachs J, Frantz S, Weismann D, Allolio B, Ertl G, Angermann CE, Stork S (2007) Complementary and incremental mortality risk prediction by cortisol and aldosterone in chronic heart failure. Circulation. 115 (13): 1754–61.https://doi.org/10.1161/CIRCULATIONAHA.106.653964

9 Girerd N, Pang PS, Swedberg K, Fought A, Kwasny MJ, Subacius H, Konstam MA, Maggioni A, Gheorghiade M, Zannad F, investigators E (2013) Serum aldosterone is associated with mortality and re-hospitalization in patients with reduced ejection fraction hospitalized for acute heart failure: analysis from the EVEREST trial. Eur J Heart Fail. 15 (11): 1228–35.https://doi.org/10.1093/eurjhf/hft100

10 Guder G, Hammer F, Deutschbein T, Brenner S, Berliner D, Deubner N, Bidlingmaier M, Ertl G, Allolio B, Angermann CE, Fassnacht M, Stork S (2015) Prognostic value of aldosterone and cortisol in patients hospitalized for acutely decompensated chronic heart failure with and without mineralocorticoid receptor antagonism. J Card Fail. 21 (3): 208–16.https://doi.org/10.1016/j.cardfail.2014.12.011

11 Volpe M (2014) Natriuretic peptides and cardio-renal disease. Int J Cardiol. 176 (3): 630–9.https://doi.org/10.1016/j.ijcard.2014.08.032

12 Volpe M, Carnovali M, Mastromarino V (2016) The natriuretic peptides system in the pathophysiology of heart failure: from molecular basis to treatment. Clin Sci (Lond). 130 (2): 57–77.https://doi.org/10.1042/CS20150469

13 Mangiafico S, Costello-Boerrigter LC, Andersen IA, Cataliotti A, Burnett JC Jr., (2013) Neutral endopeptidase inhibition and the natriuretic peptide system: an evolving strategy in cardiovascular therapeutics. Eur Heart J. 34 (12): 886–93c.https://doi.org/10.1093/eurheartj/ehs262

14 Schrier RW, Abdallah JG, Weinberger HH, Abraham WT (2000) Therapy of heart failure. Kidney Int. 57 (4): 1418–25.https://doi.org/10.1046/j.1523-1755.2000.00986.x

15 Brewster UC, Setaro JF, Perazella MA (2003) The renin-angiotensin-aldosterone system: cardiorenal effects and implications for renal and cardiovascular disease states. Am J Med Sci. 326 (1): 15–24

16 Bevan EG, Connell JM, Doyle J, Carmichael HA, Davies DL, Lorimer AR, McInnes GT (1992) Candoxatril, a neutral endopeptidase inhibitor: efficacy and tolerability in essential hypertension. J Hypertens. 10 (7): 607–13

17 Ferro CJ, Spratt JC, Haynes WG, Webb DJ (1998) Inhibition of neutral endopeptidase causes vasoconstriction of human resistance vessels in vivo. Circulation. 97 (23): 2323–30.https://doi.org/10.1161/01.CIR.97.23.2323

18 Packer M, Califf RM, Konstam MA, Krum H, McMurray JJ, Rouleau JL, Swedberg K (2002) Comparison of omapatrilat and enalapril in patients with chronic heart failure: the Omapatrilat Versus Enalapril Randomized Trial of Utility in Reducing Events (OVERTURE). Circulation. 106 (8): 920–6.https://doi.org/10.1161/01.CIR.0000029801.86489.50

19 Kostis JB, Packer M, Black HR, Schmieder R, Henry D, Levy E (2004) Omapatrilat and enalapril in patients with hypertension: the Omapatrilat Cardiovascular Treatment vs. Enalapril (OCTAVE) trial. Am J Hypertens. 17 (2): 103–11.https://doi.org/10.1016/j.amjhyper.2003.09.014

20 McMurray JJ, Packer M, Desai AS, Gong J, Lefkowitz MP, Rizkala AR, Rouleau JL, Shi VC, Solomon SD, Swedberg K, Zile MR, Investigators P-H, Committees (2014) Angiotensin-neprilysin inhibition versus enalapril in heart failure. N Engl J Med. 371 (11): 993–1004.https://doi.org/10.1056/NEJMoa1409077

21 Langenickel TH, Dole WP (2012) Angiotensin receptor-neprilysin inhibition with LCZ696: a novel approach for the treatment of heart failure.. Drug Discovery Today: Therapeutic Strategies. 9 (4): e131–e9.https://doi.org/10.1016/j.ddstr.2013.11.002

22 Yancy CW, Jessup M, Bozkurt B, Butler J, Casey DE Jr.,, Drazner MH, Fonarow GC, Geraci SA, Horwich T, Januzzi JL, Johnson MR, Kasper EK, Levy WC, Masoudi FA, McBride PE, McMurray JJ, Mitchell JE, Peterson PN, Riegel B, Sam F, Stevenson LW, Tang WH, Tsai EJ, Wilkoff BL, American College of Cardiology F, American Heart Association Task Force on Practice G (2013) 2013 ACCF/AHA guideline for the management of heart failure: a report of the American College of Cardiology Foundation/American Heart Association Task Force on Practice Guidelines. J Am Coll Cardiol. 62 (16): e147–239.https://doi.org/10.1016/j.jacc.2013.05.019

23 Shi J, Wang X, Nguyen J, Wu AH, Bleske BE, Zhu HJ (2016) Sacubitril Is Selectively Activated by Carboxylesterase 1 (CES1) in the Liver and the Activation Is Affected by CES1 Genetic Variation. Drug Metab Dispos. 44 (4): 554–9.https://doi.org/10.1124/dmd.115.068536

24 Solomon SD, Zile M, Pieske B, Voors A, Shah A, Kraigher-Krainer E, Shi V, Bransford T, Takeuchi M, Gong J, Lefkowitz M, Packer M, McMurray JJ, Prospective comparison of AwARBoMOhfwpefI (2012) The angiotensin receptor neprilysin inhibitor LCZ696 in heart failure with preserved ejection fraction: a phase 2 double-blind randomised controlled trial. Lancet. 380 (9851): 1387–95.https://doi.org/10.1016/S0140-6736(12)61227-6

25 Ponikowski P, Voors AA, Anker SD, Bueno H, Cleland JG, Coats AJ, Falk V, Gonzalez-Juanatey JR, Harjola VP, Jankowska EA, Jessup M, Linde C, Nihoyannopoulos P, Parissis JT, Pieske B, Riley JP, Rosano GM, Ruilope LM, Ruschitzka F, Rutten FH, van der Meer P, Authors/Task Force M, Document R (2016) 2016 ESC Guidelines for the diagnosis and treatment of acute and chronic heart failure: The Task Force for the diagnosis and treatment of acute and chronic heart failure of the European Society of Cardiology (ESC). Developed with the special contribution of the Heart Failure Association (HFA) of the ESC. Eur J Heart Fail. 18 (8): 891–975.https://doi.org/10.1002/ejhf.592

26 Yancy CW, Jessup M, Bozkurt B, Butler J, Casey DE, Jr.,, Colvin MM, Drazner MH, Filippatos G, Fonarow GC, Givertz MM, Hollenberg SM, Lindenfeld J, Masoudi FA, McBride PE, Peterson PN, Stevenson LW, Westlake C (2016) 2016 ACC/AHA/HFSA Focused Update on New Pharmacological Therapy for Heart Failure: An Update of the 2013 ACCF/AHA Guideline for the Management of Heart Failure: A Report of the American College of Cardiology/American Heart Association Task Force on Clinical Practice Guidelines and the Heart Failure Society of America. J Am Coll Cardiol. 68 (13): 1476–88.https://doi.org/10.1016/j.jacc.2016.05.011

27 Sayer MB, Atkins CE, Fujii Y, Adams AK, DeFrancesco TC, Keene BW (2009) Acute effect of pimobendan and furosemide on the circulating renin-angiotensin-aldosterone system in healthy dogs. J Vet Intern Med. 23 (5): 1003–6.https://doi.org/10.1111/j.1939-1676.2009.0367.x

28 Watkins L, Jr.,, Burton JA, Haber E, Cant JR, Smith FW, Barger AC (1976) The renin-angiotensin-aldosterone system in congestive failure in conscious dogs. J Clin Invest. 57 (6): 1606–17.https://doi.org/10.1172/JCI108431

29 Cowley AW, Jr., Guyton AC (1972) Quantification of intermediate steps in the renin-angiotensin-vasoconstrictor feedback loop in the dog. Circ Res. 30 (5): 557–66.https://doi.org/10.1161/01.RES.30.5.557

30 Guyton AC, Coleman TG, Cowley AW, Jr., Liard JF, Norman RA, Jr., Manning RD, Jr. (1972) Systems analysis of arterial pressure regulation and hypertension. Ann Biomed Eng. 1 (2): 254–81.https://doi.org/10.1007/BF02584211

31 Mochel JP, Danhof M (2015) Chronobiology and Pharmacologic Modulation of the Renin-Angiotensin-Aldosterone System in Dogs: What Have We Learned? Rev Physiol Biochem Pharmacol. 169: 43–69.https://doi.org/10.1007/112_2015_27

32 Mochel JP, Fink M, Peyrou M, Soubret A, Giraudel JM, Danhof M (2015) Pharmacokinetic/Pharmacodynamic Modeling of Renin-Angiotensin Aldosterone Biomarkers Following Angiotensin-Converting Enzyme (ACE) Inhibition Therapy with Benazepril in Dogs. Pharm Res. 32 (6): 1931–46.https://doi.org/10.1007/s11095-014-1587-9

33 Kjolby MJ, Kompanowska-Jezierska E, Wamberg S, Bie P (2005) Effects of sodium intake on plasma potassium and renin angiotensin aldosterone system in conscious dogs. Acta Physiol Scand. 184 (3): 225–34.https://doi.org/10.1111/j.1365-201X.2005.01452.x

34 Portaluppi F, Smolensky MH, Touitou Y (2010) Ethics and methods for biological rhythm research on animals and human beings. Chronobiol Int. 27 (9-10): 1911–29.https://doi.org/10.3109/07420528.2010.516381

35 Gu J, Noe A, Chandra P, Al-Fayoumi S, Ligueros-Saylan M, Sarangapani R, Maahs S, Ksander G, Rigel DF, Jeng AY, Lin TH, Zheng W, Dole WP (2010) Pharmacokinetics and pharmacodynamics of LCZ696, a novel dual-acting angiotensin receptor-neprilysin inhibitor (ARNi). J Clin Pharmacol. 50 (4): 401–14.https://doi.org/10.1177/0091270009343932

36 King JN, Mauron C, Kaiser G (1995) Pharmacokinetics of the active metabolite of benazepril, benazeprilat, and inhibition of plasma angiotensin-converting enzyme activity after single and repeated administrations to dogs. Am J Vet Res. 56 (12): 1620–8

37 Muller AF, Manning EL, Riondel AM (1958) Influence of position and activity on the secretion of aldosterone. Lancet. 1 (7023): 711–3.https://doi.org/10.1016/S0140-6736(58)91137-1

38 Mochel JP, Fink M, Peyrou M, Desevaux C, Deurinck M, Giraudel JM, Danhof M (2013) Chronobiology of the renin-angiotensin-aldosterone system in dogs: relation to blood pressure and renal physiology. Chronobiol Int. 30 (9): 1144–59.https://doi.org/10.3109/07420528.2013.807275

39 Mochel JP, Fink M, Bon C, Peyrou M, Bieth B, Desevaux C, Deurinck M, Giraudel JM, Danhof M (2014) Influence of feeding schedules on the chronobiology of renin activity, urinary electrolytes and blood pressure in dogs. Chronobiol Int. 31 (5): 715–30.https://doi.org/10.3109/07420528.2014.897711

40 King JN, Maurer M, Morrison CA, Mauron C, Kaiser G (1997) Pharmacokinetics of the angiotensin-converting-enzyme inhibitor, benazepril, and its active metabolite, benazeprilat, in dog. Xenobiotica. 27 (8): 819–29.https://doi.org/10.1080/004982597240181

41 Zhdanova IV, Simmons M, Marcus JN, Busza AC, Leclair OU, Taylor JA (1999) Nocturnal increase in plasma cGMP levels in humans. J Biol Rhythms. 14 (4): 307–13.https://doi.org/10.1177/074873099129000722

42 Azizi M, Menard J (2004) Combined blockade of the renin-angiotensin system with angiotensin-converting enzyme inhibitors and angiotensin II type 1 receptor antagonists. Circulation. 109 (21): 2492–9.https://doi.org/10.1161/01.CIR.0000131449.94713.AD

43 Trivedi RK, Polhemus DJ, Li Z, Yoo D, Koiwaya H, Scarborough A, Goodchild TT, Lefer DJ (2018) Combined Angiotensin Receptor-Neprilysin Inhibitors Improve Cardiac and Vascular Function Via Increased NO Bioavailability in Heart Failure. J Am Heart Assoc. 7 (5).https://doi.org/10.1161/JAHA.117.008268

44 Kobalava Z, Kotovskaya Y, Averkov O, Pavlikova E, Moiseev V, Albrecht D, Chandra P, Ayalasomayajula S, Prescott MF, Pal P, Langenickel TH, Jordaan P, Rajman I (2016) Pharmacodynamic and Pharmacokinetic Profiles of Sacubitril/Valsartan (LCZ696) in Patients with Heart Failure and Reduced Ejection Fraction. Cardiovasc Ther. 34 (4): 191–8.https://doi.org/10.1111/1755-5922.12183

45 Packer M, McMurray JJ, Desai AS, Gong J, Lefkowitz MP, Rizkala AR, Rouleau JL, Shi VC, Solomon SD, Swedberg K, Zile M, Andersen K, Arango JL, Arnold JM, Belohlavek J, Bohm M, Boytsov S, Burgess LJ, Cabrera W, Calvo C, Chen CH, Dukat A, Duarte YC, Erglis A, Fu M, Gomez E, Gonzalez-Medina A, Hagege AA, Huang J, Katova T, Kiatchoosakun S, Kim KS, Kozan O, Llamas EB, Martinez F, Merkely B, Mendoza I, Mosterd A, Negrusz-Kawecka M, Peuhkurinen K, Ramires FJ, Refsgaard J, Rosenthal A, Senni M, Sibulo AS, Jr., Silva-Cardoso J, Squire IB, Starling RC, Teerlink JR, Vanhaecke J, Vinereanu D, Wong RC, Investigators P-H, Coordinatorsdagger (2015) Angiotensin receptor neprilysin inhibition compared with enalapril on the risk of clinical progression in surviving patients with heart failure. Circulation. 131 (1): 54–61.https://doi.org/10.1161/CIRCULATIONAHA.114.013748

46 Johns DW, Peach MJ, Gomez RA, Inagami T, Carey RM (1990) Angiotensin II regulates renin gene expression. Am J Physiol. 259 (6 Pt 2): F882–7.https://doi.org/10.1152/ajprenal.1990.259.6.F882

47 Keeton TK, Campbell WB (1980) The pharmacologic alteration of renin release. Pharmacol Rev. 32 (2): 81–227

48 Kurtz A, Della Bruna R, Pfeilschifter J, Taugner R, Bauer C (1986) Atrial natriuretic peptide inhibits renin release from juxtaglomerular cells by a cGMP-mediated process. Proc Natl Acad Sci U S A. 83 (13): 4769–73.https://doi.org/10.1073/pnas.83.13.4769

49 Erdos EG, Skidgel RA (1989) Neutral endopeptidase 24.11 (enkephalinase) and related regulators of peptide hormones. FASEB J. 3 (2): 145–51.https://doi.org/10.1096/fasebj.3.2.2521610

50 von Lueder TG, Atar D, Krum H (2014) Current role of neprilysin inhibitors in hypertension and heart failure. Pharmacol Ther. 144 (1): 41–9.https://doi.org/10.1016/j.pharmthera.2014.05.002

51 van de Wal RM, Plokker HW, Lok DJ, Boomsma F, van der Horst FA, van Veldhuisen DJ, van Gilst WH, Voors AA (2006) Determinants of increased angiotensin II levels in severe chronic heart failure patients despite ACE inhibition. Int J Cardiol. 106 (3): 367–72.https://doi.org/10.1016/j.ijcard.2005.02.016

52 MacFadyen RJ, Lee AF, Morton JJ, Pringle SD, Struthers AD (1999) How often are angiotensin II and aldosterone concentrations raised during chronic ACE inhibitor treatment in cardiac failure? Heart. 82 (1): 57–61.https://doi.org/10.1136/hrt.82.1.57

53 Rocha R, Chander PN, Zuckerman A, Stier CT, Jr. (1999) Role of aldosterone in renal vascular injury in stroke-prone hypertensive rats. Hypertension. 33 (1 Pt 2): 232–7.https://doi.org/10.1161/01.HYP.33.1.232

54 Latini R, Masson S, Anand I, Salio M, Hester A, Judd D, Barlera S, Maggioni AP, Tognoni G, Cohn JN, Val-He FTI (2004) The comparative prognostic value of plasma neurohormones at baseline in patients with heart failure enrolled in Val-HeFT. Eur Heart J. 25 (4): 292–9.https://doi.org/10.1016/j.ehj.2003.10.030

55 Roig E, Perez-Villa F, Morales M, Jimenez W, Orus J, Heras M, Sanz G (2000) Clinical implications of increased plasma angiotensin II despite ACE inhibitor therapy in patients with congestive heart failure. Eur Heart J. 21 (1): 53–7.https://doi.org/10.1053/euhj.1999.1740

56 Swedberg K, Eneroth P, Kjekshus J, Wilhelmsen L (1990) Hormones regulating cardiovascular function in patients with severe congestive heart failure and their relation to mortality. CONSENSUS Trial Study Group. Circulation. 82 (5): 1730–6.https://doi.org/10.1161/01.CIR.82.5.1730

57 Jordaan P (2011) Changes in RAAS (renin angiotensin aldosterone system) biomarkers in patients with stable chronic heart failure following short-term angiotensin receptor neprilysin inhibitor (ARNI) treatment. SA Heart. 8: 236

58 Franco-Saenz R, Atarashi K, Takagi M, Takagi M, Mulrow PJ (1989) Effect of atrial natriuretic factor on renin and aldosterone. J Cardiovasc Pharmacol. 13 Suppl 6: S31–5

59 Mochel JP, Fink M (2012) Response to letter from Atkins et al. J Vet Pharmacol Ther. 35 (5): 516–8

